# Diverse viruses in deep-sea hydrothermal vent fluids have restricted dispersal across ocean basins

**DOI:** 10.1101/2020.06.04.125666

**Authors:** Elaina Thomas, Rika Anderson, Viola Li, Jenni Rogan, Julie A. Huber

## Abstract

In the ocean, viruses impact microbial mortality, regulate biogeochemical cycling, and alter the metabolic potential of microbial lineages. At deep-sea hydrothermal vents, abundant viruses infect a wide range of hosts among the archaea and bacteria that inhabit these dynamic habitats. However, little is known about viral diversity, host range, and biogeography across different vent ecosystems, which has important implications for how viruses manipulate microbial function and evolution. Here, we examined viral diversity, viral and host distribution, and viral-host interactions in microbial metagenomes generated from venting fluids from several vent sites within three different geochemically and geographically distinct hydrothermal systems: Piccard and Von Damm vent fields at the Mid-Cayman Rise in the Caribbean Sea, and at several vent sites within Axial Seamount in the Pacific Ocean. Analysis of viral sequences and Clustered Regularly InterSpaced Palindromic Repeats (CRISPR) spacers revealed highly diverse viral assemblages and evidence of active infection. Network analysis revealed that viral host range was relatively narrow, with very few viruses infecting multiple microbial lineages. Viruses were largely endemic to individual vent sites, indicating restricted dispersal, and in some cases viral assemblages persisted over time. Thus, we show that hydrothermal vent fluids are home to novel, diverse viral assemblages that are highly localized to specific regions and taxa.

**Importance:** Viruses play important roles in manipulating microbial communities and their evolution in the ocean, yet not much is known about viruses in deep-sea hydrothermal vents. However, viral ecology and evolution are of particular interest in hydrothermal vent habitats because of their unique nature: previous studies have indicated that most viruses in hydrothermal vents are temperate rather than lytic, and it has been established that rates of horizontal gene transfer (HGT) are particularly high among thermophilic vent microbes, and viruses are common vectors for HGT. If viruses have broad host range or are widespread across vent sites, they have increased potential to act as gene-sharing “highways” between vent sites. By examining viral diversity, distribution, and infection networks across disparate vent sites, this study provides the opportunity to better characterize and constrain the viral impact on hydrothermal vent microbial communities. We show that viruses in hydrothermal vents are diverse and apparently active, but most have restricted host range and are not widely distributed among vent sites. Thus, the impacts of viral infection are likely to be highly localized and constrained to specific taxa in these habitats.

## Introduction

Deep-sea hydrothermal vents are regions on the seafloor where high-temperature hydrothermal vent fluid is created from water-rock reactions deep within the oceanic crust, mixing with seawater beneath and at the seafloor to create a dynamic, gradient-dominated habitat that supports diverse microbial communities. These low-temperature diffuse vent fluids are hotspots of primary production in the deep ocean, dominated by chemolithoautotrophic bacteria and archaea carrying out a variety of metabolisms utilizing hydrogen, sulfur compounds, nitrate, and methane (1–7). An important, though understudied, driver of microbial diversity and evolution in hydrothermal systems is viruses. Viruses are major sources of microbial mortality in marine systems and play key roles in mediating biogeochemical cycling and shaping microbial community structure (8–11). At deep-sea hydrothermal vents, viruses are abundant (12) and infect a wide range of microbial hosts (13). Temperate viruses, which often integrate into the genomes of their microbial hosts, are particularly abundant in hydrothermal fluids compared to other marine habitats (14, 15). The high abundance of temperate viruses suggests that there are unique attributes to the vent environment that influence viral infection strategies. Viruses can “metabolically reprogram” their microbial hosts via the introduction and expression of auxiliary metabolic genes (AMGs) (16), which has profound impacts on the ecology and evolution of microbial populations. Like horizontally transferred genes, integrated prophage with AMGs can alter the functional potential of a given organism, allowing it to adapt to changing conditions or expand to new ecological niches. Viruses in hydrothermal habitats have been found to encode AMGs (17–19), and thus can manipulate the metabolic potential of microbial populations in hydrothermal vents.

Viruses also function as vectors of horizontal gene transfer via transduction. Previous work has suggested that horizontal gene transfer is particularly prevalent among microbes inhabiting high-temperature environments (20–22). Transposases, which catalyze the movement of mobile genetic elements within and between genomes (23, 24), are abundant in hydrothermal vent sites (18, 25). Given the abundance of viruses and the inferred high rates of transduction at deep-sea vents, it is likely that viruses are an important vector for horizontal gene transfer in these dynamic systems. Therefore, a clear understanding of which viruses infect which hosts, and what genes those viruses carry, can provide insight into highways of gene sharing in the deep sea.

One large gap in our understanding of eco-evolutionary dynamics in deep-sea hydrothermal vents is how broadly viruses are distributed within and between vent systems, and how viral interactions shift across hydrothermal vent types. Previous work has shown that microbial communities exhibit high endemism locally (4, 7, 26, 27), but some cosmopolitan species are distributed globally (28–30). However, less is known about the geographic distribution of viral populations. If viruses exhibit a restricted distribution, this would limit their role as vectors of gene flow across and between hydrothermal systems. Microbial populations and their viruses are also limited by environmental conditions: hydrothermal systems hosted in basalt rocks are characterized by metal-enriched, low pH fluids up to 400°C (31). In contrast, hydrothermal systems hosted in peridotite are influenced by serpentinization and feature organic carbon-enriched, high pH fluids with slightly lower temperature (32). Microbial populations in basalt-hosted and peridotite-hosted vents exhibit distinct patterns of genomic variation (33), suggesting that microbes are subject to different selection pressures depending on the vent type, but we know little about the role viruses play in driving those differences, nor how viral diversity varies across hydrothermal systems.

To gain further insight into viral diversity, distribution, and host range across hydrothermal systems, we compared viral sequences recovered from microbial metagenomes collected from two hydrothermal regions in two distinct ocean basins: Axial Seamount, a submarine volcano located on the Juan de Fuca Ridge in the Pacific Ocean at ~1520 meters depth, and the Mid-Cayman Rise, an ultraslow spreading ridge in the Caribbean Sea. Axial Seamount is a basalt-hosted, magma-driven system with fluids that are low in pH and high in carbon dioxide and hydrogen sulfide (34). Microbial communities at Axial are spatially restricted but temporally stable at individual vent sites over time (4, 27). In contrast, the Mid-Cayman Rise hosts two geologically and geochemically distinct hydrothermal vent fields in close proximity to each other: Piccard hydrothermal field, located in basalt rocks along the ridge axis, is the deepest hydrothermal vent field discovered to date (~4950 meters depth) and is characterized by fluids that are acidic with high hydrogen and hydrogen sulfide content, whereas the Von Damm vent field, located approximately 20 km away on a nearby massif at ~2350 meters depth, is influenced by serpentinization and is characterized by fluids that are high in hydrogen, methane, and small carbon compounds (35–38). Previous work found distinct community composition but similar functional potential in the microbial communities at Piccard and Von Damm (7, 27, 33, 39, 40). Here, we compare viral sequences in venting fluids from Von Damm vent field, Piccard vent field, and Axial Seamount in a comparative survey of viral diversity and gene content across geographically and geochemically distinct hydrothermal vent sites. By examining viruses from the microbial fraction (0.22 μm), we are closely examining the relationship between prophage, actively infecting viruses, and their hosts. We examined viral sequences identified in the metagenomes as well as CRISPR sequences, which provide context for historical viral infections within each of the communities, and provide a means to match viruses with putative hosts. We show that viral populations have high diversity but restricted distribution and host range across hydrothermal systems, limiting the capacity of viruses to act as vectors of gene flow between disparate hosts and vent locations.

## Methods

### Sample collection and DNA preparation and sequencing

Low-temperature diffuse flow fluid samples were collected from the vent fields Piccard and Von Damm at the Mid-Cayman Rise in January 2012 and June 2013 during research cruises on the R/V Atlantis and R/V Falkor, respectively. We analyzed a total of 11 metagenomes from 8 different vents at Von Damm vent field and 4 metagenomes from 4 different vents at Piccard vent field. Sample locations, depth, and metagenomic data are described in Supplementary Table 1. The ROV Jason II and Mat Sampler were used to collect the 2012 Mid-Cayman Rise samples, as previously described in (41). The 2013 Mid-Cayman Rise samples were collected using the SUPR version 2 sampler and HROV Nereus (42). For microbial DNA collection, approximately 3 to 6 L of diffuse flow fluid were pumped through 0.22 μm Sterivex filters (Millipore). Shipboard, the filters were flooded with RNALater (Ambion), sealed with Luer Caps, stored in sterile Falcon tubes, and frozen at −80 °C. Sample collection and preservation are further described in Anderson et al. (33) and Reveillaud et al. (40). Genomic DNA was extracted and metagenomic libraries constructed as described in Anderson et al. (33). Sequencing was done on an Illumina Hi Seq 1000 at the W.M. Keck Facility in the Josephine Bay Paul Center at the Marine Biological Laboratory.

Diffuse flow fluid samples were collected from Axial Seamount in September 2013, August 2014, and August 2015 (approximately five months after the eruption of Axial Seamount) during research cruises on the R/V Falkor and R/V Thompson in 2013, R/V Brown in 2014, and R/V Thompson in 2015 (Supplementary Table 1). Diffuse flow samples were collected from four different vent fields within Axial Seamount (ASHES, International District, North Rift Zone, and Dependable) using the ROVs ROPOS and Jason. We analyzed a total of 16 metagenomes from 10 different vents, 1 plume, and 2 deep seawater samples at Axial Seamount. For collection of microbial DNA, 3 L of diffuse fluid was pumped through 0.22 μm, 47 mm GWSP filters (Millipore) and the filters were flooded with RNALater (Ambion) on the seafloor, as described in Fortunato et al. (4). Fluids from a hydrothermal plume above Anemone and background seawater were collected using a Seabird SBE911 CTD and 10 L Niskin bottles and 3 L of the plume and seawater fluid was filtered through 0.22 μm Sterivex filters (Millipore). DNA was extracted from the filters using a phenol-chloroform method adapted from Crump et al. (43) and Zhou et al. (44). The Ovation Ultralow Library DR multiplex system (Nugen) was used to prepare metagenomic libraries following the manufacturer instructions. DNA extraction and metagenomic library construction is further described in Fortunato and Huber (45). The 2013 and 2014 samples were sequenced on an Illumina HiSeq 1000 and the 2015 samples on a NextSeq 500. Sequencing was done at the W.M. Keck sequencing facility at the Marine Biological Laboratory.

For all metagenomes, paired-end partially overlapping reads were merged and quality filtered using the illumina-utils package (46) using the iu-quality-filter-minoche flag, then assembled using idba-ud (47) v1.1.2 with default settings. All data from both sites is available in the European Nucleotide Archives archives under study accession number PRJEB15541 for the Mid-Cayman Rise and under study accession numbers PRJEB7866, PRJEB12000, and PRJEB19456 for Axial samples in the years 2013, 2014 and 2015 respectively (Supplementary Table 1).

### Identification of CRISPR loci, viral populations, and spacer assemblages

Crass (48) v1.0.1 was used to identify CRISPR loci (i.e., unique direct repeat types) and CRISPR spacers in the metagenomic reads. Viral contigs in the metagenomic assemblies were identified with VirFinder (48, 49) v1.1 using a p-value threshold of 0.05. Given that viruses from hydrothermal systems are not well-represented in sequence databases, we elected to use VirFinder for identification of viral contigs for diversity and abundance analysis because VirFinder is a k-mer frequency-based method that avoids gene-based similarity searches, and thus has higher potential to identify novel viruses. We used ClusterGenomes (https://bitbucket.org/MAVERICLab/stampede-clustergenomes) v1.1.3 (95% identity, 80% coverage) to identify viral clusters (or viral operational taxonomic units, vOTUs) of viral contigs. To cluster the CRISPR spacers, we performed an all-v-all blast using BLASTn v2.5.0 (E-value threshold of 10^-08^) and then clustered using Markov Cluster Algorithm (MCL) (50) v14-137 (inflation 1.2 and scheme 7) based on bitscore.

### Recovery of MAGs

All metagenome assembled genomes (MAGs) were recovered from metagenomic assemblies using anvi’o (51). Supervised clustering was used to recover bins from Piccard and Von Damm contigs using anvi’o v2.1.0 (33). MAGs from Axial Seamount assemblies were recovered using unsupervised binning with CONCOCT (50, 52, 53) within the anvi’o v4.0 pipeline, followed by manual refinement within anvi’o (39). For all analyses, only bins with >=70% completeness and <=10% redundancy were retained as MAGs for this analysis. The MAGs were assigned taxonomies using PhyloSift (50, 52) v1.0.1 as described in Anderson et al. (33). Only MAGs for which taxonomies could be identified were used in analyses. MAG coverage was normalized by the number of merged reads in the metagenome. All of the Mid-Cayman Rise MAGs were previously described in Anderson et al. (33).

### Taxonomic identification of microbial and viral reads

The reads from all metagenomes were mapped to the Silva SSU and LSU Parc databases (54, 55) (release 132) with bowtie2 (56) v2.2.9 using default settings and local alignment. Mapped reads were assigned taxonomies using the classify.seqs function in mothur (57, 58) v1.38.1 with the Silva 16S rRNA database (release 119) and a cutoff of 50. Reads mapping to 16S rRNA gene sequences that were classified as eukaryotes were excluded from analyses.

In order to identify the viral taxa at Von Damm, Piccard, and Axial Seamount, we conducted a blast of the representative sequence of each vOTU against all of the viral genomic sequences in RefSeq (downloaded on September 16th, 2020; E-value <= 10-5). The best match for each vOTU representative sequence was selected based on E-value and percent identity. The taxonomy of the RefSeq viral sequence with the best alignment was assigned to the given vOTU. To quantify the relative abundance of viral families in each sample, the reads that mapped to vOTUs of a viral family were divided by the total number of reads in the sample.

### Viral, spacer, and host diversity

Rarefaction curves for vOTUs, CRISPR spacer clusters, and reads matching 16S rRNA genes categorized at the class level were created using the Vegan R package (59) v2.4-5. The number of vOTUs per contig and spacer clusters per read (paired reads) were used as proxies for viral diversity, and the number of different taxa at the class level matching 16S rRNA genes reflected microbial diversity, calculated on a per read basis (merged reads).

### Viral, spacer, and CRISPR relative abundance

To calculate relative viral abundance, the reads in each of the metagenomes were mapped against all of the viral contigs from the corresponding geographic region using bowtie2 (56) v2.2.9. The reads from each sample were mapped against all of the viral contigs from the corresponding geographic region rather than solely the viral contigs in the sample because there were viral reads in samples that did not assemble into viral contigs. This method therefore allowed the identification of more viral sequences. The number of reads that mapped to viral contigs was normalized by the number of merged reads as a measure of relative viral abundance. This measure of relative viral abundance reflects only the proportion of viruses that were retained on the filter as viral capsids or prophages. We used the number of spacers per read and CRISPR direct repeat types per read as measures of spacer and CRISPR relative abundance, respectively. Paired rather than merged reads were used for these analyses. Relative abundance and diversity of viruses, microbes, CRISPR spacers and CRISPR loci were visualized using the Seaborn library within Python (60).

### Relative compositions and abundances of viral populations, spacer assemblages, and hosts

The relative compositions of the microbial community, the viral assemblage, and CRISPR spacers were compared between vent sites within Von Damm, Piccard, and Axial Seamount. To calculate the relative abundance of each vOTU in each sample, the number of reads in the sample that mapped to the viral contigs in the vOTU was determined using bowtie2 (56) v2.2.9. The number of reads in each metagenome that mapped to the vOTU was normalized by the total length of the viral contigs in the vOTU and the number of merged reads in the metagenome. We defined the most common vOTUs as the six clusters with the highest relative abundance in each sample. The relative abundance of each CRISPR spacer cluster in each sample was calculated as the percent of spacers in the sample that were part of the spacer cluster. For spacer clusters, we defined the most common as the three clusters with the highest relative abundance in each sample. To compare the compositions of viral contigs and CRISPR spacers, all of the spacers were blasted against all of the viral contigs using blastn (E-value <=10^-05^, <=1 mismatch, as per Emerson et al. (61). The relative abundance of each microbial host was measured as the number of reads that mapped to 16S rRNA gene sequences of the given taxon, normalized by the number of 16S rRNA gene reads in the sample. Analyses of microbial taxa were done at either the class level or the lowest taxonomic level available, depending on the analysis.

Microbiome datasets are compositional in nature because sequencing instruments impose an arbitrary total (62). Therefore, to conduct hierarchical clustering of samples based on viral, spacer, and host composition, we used the protocol outlined by Gloor et al. (62) and Gloor and Reid (63) for computing distances between samples containing compositional data. For hierarchical clustering, we did not normalize; we performed analyses on the number of reads that mapped to each vOTU, the number of CRISPR spacers in each spacer cluster, and the number of reads that mapped to 16S rRNA gene sequences of each microbial host in each sample; for microbial hosts, we did not include reads that mapped to unclassified sequences or sequences classified as eukaryotes. We replaced zero counts with estimates using the count zero multiplicative method for vOTUs and hosts and the Bayes-Laplace Bayesian multiplicative method for spacer clusters via the zCompositions R package (64) v1.2.0. Using the CoDaSeq R package (https://github.com/ggloor/CoDaSeq) (62) v0.99.3, we applied a centered log-ratio (clr) transformation to the count data (that lacked zero counts), thereby capturing the ratios between parts. To calculate distances between samples for hierarchical clustering, we used the ward.D2 method on the transformed counts.

### Networks of viral infection

Infection networks of viral-host interactions were created using CRISPR sequences to connect microbial hosts with clusters of viral contigs, adapted from the methods used by Daly et al. (65) and Emerson et al. (61). First, MAGs were connected to CRISPR direct repeat types within each sample using BLASTn v2.5.0 (E-value <=10^-10^, 100% nucleotide identity, as per Emerson et al. (61). Then, each CRISPR direct repeat type was matched to a set of CRISPR spacers as identified by Crass (Skennerton et al. 2013) v1.0.1. Finally, the CRISPR spacers in each sample were matched to viral contigs in the corresponding region using BLASTn v2.5.0 with an E-value cutoff of 10^-05^ and a maximum of one mismatch, as per Emerson et al. (61). Only one mismatch was allowed because resistance has been found to be lost by single nucleotide differences between bacterial spacers and target phage sequences (66). In contrast, resistance by archaeal CRISPR systems can still be provided when there are up to three mismatches between spacers and target phage sequences (67, 68). We did not allow for more mismatches between archaeal spacers and phage sequences because resistance is weakened by more mismatches (68). Each direct repeat type provided by Crass v1.0.1 does not necessarily represent an individual CRISPR locus (48).

### Data analysis software

The majority of analyses were conducted in RStudio (69) v1.0.136. R packages used were readr (70) v1.1.1, readxl (71) v1.0.0, tidyr (72) v0.7.2, stringr (73) v1.3.0, dplyr (74) v0.7.4, ggnetwork (75) v0.5.1, statnet (76) v2016.9, ggpubr (77) v0.1.7, ggplot2 (78) v2.2.1, and svglite (79) v1.2.1.

## Results

### Identification of putative viral contigs

We used VirFinder to identify putative viral contigs because it is a k-mer frequency-based method that has higher potential to identify novel viruses (Ren *et al*., 2017). However, this method allows for the possibility of false positives. VirFinder assigns scores between 0-1 to indicate the likelihood that a contig is viral, with higher values reflecting a higher likelihood that the sequence is viral. Almost all VirFinder sequences from Axial Seamount (99.96%), Piccard (98.54%), and Von Damm (98.96%) were assigned scores greater than 0.7. 47.35% of Axial Seamount, 44.21% of Piccard, and 37.92% of Von Damm VirFinder sequences had scores greater than 0.9. The mean score of VirFinder sequences was 0.89, 0.88 and 0.87 for Axial Seamount, Piccard, and Von Damm, respectively. Supplementary Figure 1a shows the distribution of scores for the contigs identified with VirFinder at the Axial Seamount, Piccard, and Von Damm vent fields. Similarly, the majority of contigs identified as putatively viral were short, with shorter average contigs at Axial Seamount (625 bp) than those at Piccard (2172 bp) or Von Damm (2037 bp). The VirFinder score did not increase with sequence length (Supplementary Figure 1b).

### Taxonomy of viruses and hosts

All analyses were carried out from diffuse flow fluids sampled directly from the vent orifice, as well as vent plume waters, which were sampled up to 100m above the vent orifice and had much lower temperatures (Supplementary Table 1). It is possible to examine viruses in 0.22 μm-filtered fluids because this fraction includes integrated prophage, lytic infections in progress, and free viral particles captured on filters. However, our analysis misses free viral particles that were not retained on the filters, and we are only capturing the viral diversity and variation across sites based on the viral sequences identified in the metagenomes.

The viral sequences identified at Axial Seamount, Von Damm, and Piccard vents represent 28 families of viruses (Figure 1). 13 of the viral families were present at all three vent fields. 14 viral families were present at both of the Mid-Cayman Rise vent fields, Von Damm and Piccard, while Axial Seamount shared 13 viral families with each of the Mid-Cayman Rise vent fields. We observed a distinct difference in viral taxonomic groups between the vent fields: whereas the *Myoviridae* viral family had the highest relative abundance in 12 of the 16 Axial Seamount samples, the *Myoviridae* were in much lower abundance at Von Damm and Piccard, which had higher abundances of reads matching the *Podoviridae* family of viruses and the *Guttaviridae*.

**Figure 1.**
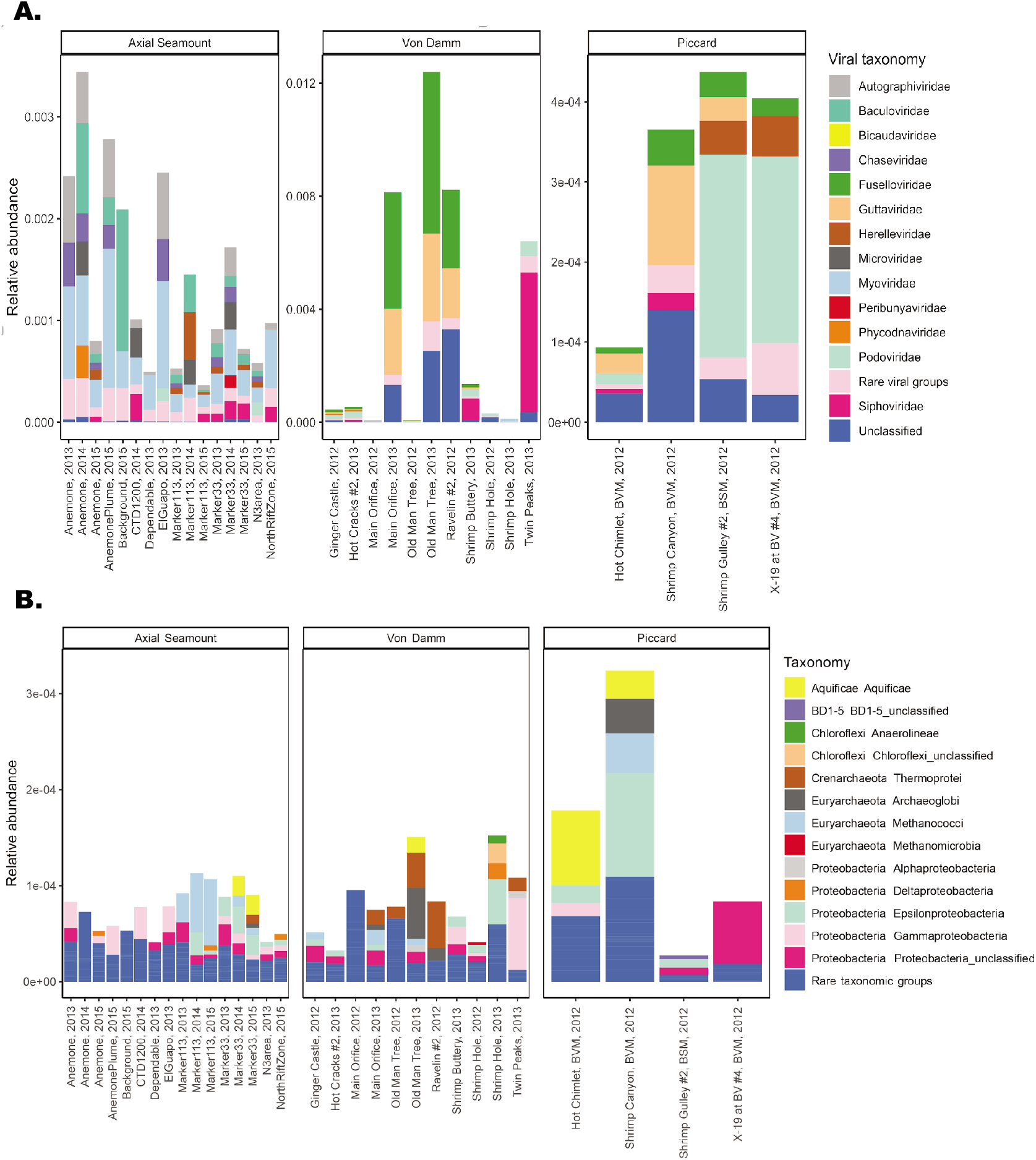
Taxonomy of A) viral and B) microbial reads in the Von Damm, Piccard, and Axial Seamount vent fields. In A) the y-axis represents the number of reads that mapped to VirFinder contigs of each viral taxonomy normalized by the number of reads in each sample. Rare viral groups represent viral taxonomies that comprised less than 5% of reads that mapped to viral contigs in a given sample. In B) the y-axis represents the number of reads that mapped to 16S rRNA gene sequences of each host normalized by the number of reads in each sample. Reads that mapped to 16S gene rRNA sequences that were unclassified or classified as eukaryotes were excluded. Rare taxonomic groups represent taxonomies that comprised less than 1% of reads that mapped to 16S rRNA gene sequences in a given sample.

### Relative abundance of viral and microbial reads

In order to examine the distribution and abundance of viruses across individual vent sites, vent fields, and regions, we quantified the relative abundance of viral sequences, CRISPRs, and spacers in each of the metagenomes (Figure 2, samples from background seawater and plumes are excluded; Supplementary Table 2). We also quantified the diversity and abundance of loci and spacers within CRISPRs each of the metagenomes. CRISPR loci are a microbial immune system found in archaeal and bacterial genomes, consisting of direct repeats interspersed by “spacers,” which match foreign DNA (predominantly viruses, but also including plasmids and other forms of foreign DNA) that the cell has been exposed to previously. The relative abundance of CRISPR loci serves as an indication of how many microbial lineages use CRISPR as a mechanism for viral immunity. It is also important to note that CRISPR loci vary in number across microbial genomes, and we did not distinguish between CRISPR loci with the same direct repeat type. Therefore, our measure of CRISPR relative abundance is not a direct proxy for the abundance of viruses. Instead, it gives an indication of how commonly CRISPR loci are used as an antiviral mechanism within the community. Moreover, CRISPRs serve as a record of past infections, and thus while viral diversity reflects the diversity of viral particles sequenced at the time of sampling, CRISPR spacer diversity reflects the diversity of past viral infections.

**Figure 2.**
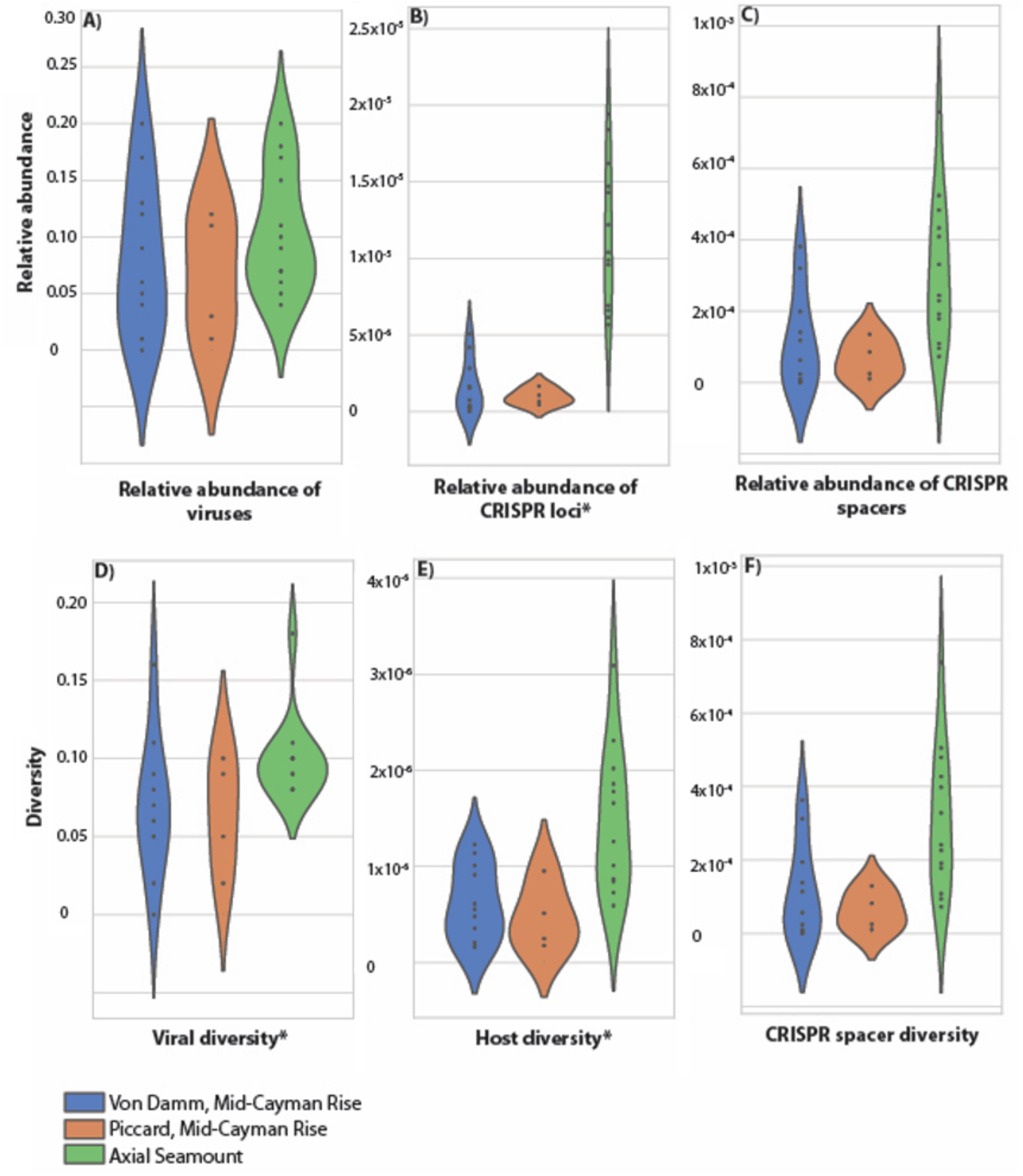
Abundance and diversity of viral sequences, CRISPR spacers, CRISPR loci, and microbes in diffuse flow samples from the Von Damm vent field (blue), Piccard vent field (orange), and Axial Seamount (green). Relative abundance reflects the relative number of reads mapping to viral contigs within each region, normalized by read abundance (see Methods). Diversity reflects the number of clusters per read, normalized by read abundance (see Methods). Note that the y-axis values are different for each variable. Values for individual samples are indicated with black dots. Violins represent the kernel density estimation of the underlying data distribution. Variables with significant differences between Von Damm, Piccard, and Axial Seamount are indicated with asterisks (based on ANOVA test). For these comparisons, samples from background seawater and plumes were excluded.

Our results indicate that the relative abundance of viral sequences (Figure 2A) was similar both across and within each of the hydrothermal vent regions we studied (p = 0.42, t-test). There was a higher relative abundance of CRISPR loci in samples collected at Axial Seamount compared to both Piccard and Von Damm vent fields (Figure 2B; p = 1.49e-05, t-test). However, there was no difference in the relative abundance of spacers within CRISPR loci per read (Figure 2C; p = 0.10, t-test). Within the Mid-Cayman Rise, we compared samples from mafic-hosted (Piccard) versus ultramafic-hosted (Von Damm) hydrothermal systems. We did not observe significant differences in the relative abundance of viral sequences, CRISPR loci, or CRISPR spacers between Piccard and Von Damm (viral sequences: p = 0.66, t-test, CRISPR loci: 0.40, t-test, spacers: 0.37, t-test) (Figure 2A-C). Similar results emerged from comparisons among vent fields within Axial Seamount: we did not observe significant differences in the relative abundance of viral sequences, CRISPR loci, or CRISPR spacers between vent fields at Axial Seamount (viral sequences: p = 0.17, t-test, CRISPR loci: 0.41, t-test, spacers: 0.11, t-test). Finally, within Axial Seamount, we also compared the relative abundance of viruses in samples taken from plume and diffuse flow hydrothermal fluid at Anemone vent. The Anemone diffuse flow samples had a higher relative abundance of CRISPR spacers and CRISPR loci compared to the Anemone plume sample, which was sampled above the vent. However, the relative abundance of viral sequences did not differ between the Anemone plume and diffuse flow samples (Supplementary Table 2).

### Diversity of viral assemblage and microbial community

Overall, viral diversity analyses revealed that the viral assemblages within these vents had high richness and were not dominated by specific viral strains. With the exception of a few samples, the rarefaction curves for the viruses and microbes did not reach saturation (Supplementary Fig. 2), indicating that the vOTUs recovered in this study did not capture the total diversity in the samples. In each sample, there were no dominant viral or spacer clusters, and each of the viral and spacer clusters present was relatively rare (Supplementary Figs. 3–4). Moreover, the viral and spacer clusters with the highest coverage did not correspond across samples: in both Piccard and Axial Seamount, one of the most common spacer clusters matched with one of the most common vOTUs by BLAST, while none of the most common spacer clusters and vOTUs aligned in Von Damm. Thus, there were no dominant viral sequences that were consistently found across all samples.

We observed a more diverse viral assemblage at Axial Seamount compared to the Mid-Cayman Rise. We observed higher richness of both viruses (Figure 2D; p = 0.015, t-test) and their microbial hosts (Figure 2E; p = 0.0056, t-test) in samples from vents at Axial Seamount compared to samples from Von Damm and Piccard vent fields. However, we did not observe meaningful correlations between vOTU, spacer cluster, or host diversity. Moreover, we did not observe significant differences in either viral or host diversity between the Piccard and Von Damm vent fields at the Mid-Cayman Rise (viral diversity: p = 0.053, t-test, host diversity: p = 0.66, t-test). We also examined the relative diversity of CRISPR spacers, which provides an indication of past viral infections rather than the current virus pool. We did not observe a significant difference in the diversity of CRISPR spacers between samples from Piccard, Von Damm, and Axial Seamount (Figure 2F, p = 0.093, t-test).

We observed significant differences in viral diversity between vent sites within Axial Seamount (p = 0.022, t-test). However, no significant differences emerged in terms of the diversity of microbial hosts and CRISPR spacers (host diversity: p = 0.74, t-test, spacer diversity: p = 0.11, t-test). At Anemone vent, the diffuse flow samples had higher CRISPR spacer and microbial diversity than the plume sample, but viral diversity did not differ between these samples (Supplementary Table 2).

### Viral distribution across vent sites

In order to characterize viral distribution and community similarity across hydrothermal vent fluids, we evaluated the extent to which viral sequences and CRISPR spacers were distributed across samples, then compared these results to the host microbial community. As before, we clustered sequences based on similarity to compare across all diffuse flow samples, excluding background seawater and plume samples. On the whole, viral sequences and CRISPR spacers had fairly limited distributions. Only a few vOTUs (0.02%) were present at all vent sites (Von Damm, Piccard, and Axial Seamount), and no CRISPR spacer clusters were present in all three regions (Table 1). The most cosmopolitan viruses and CRISPR spacers did not match each other: at Von Damm, two of the 100 most widely distributed vOTUs matched with three of the 100 most widely distributed CRISPR spacer clusters according to BLAST. At Piccard, one of the most widely distributed vOTUs aligned to one of the most widely distributed spacer clusters. At Axial Seamount, one of the 100 most widely distributed vOTUs matched with two of the 100 most widely distributed CRISPR spacer clusters. In contrast, microbial taxa were much more cosmopolitan, with ~17% of taxa shared between Von Damm, Piccard, and Axial Seamount according to 16S rRNA gene classification at the lowest taxonomic level available (Table 1). Viral and CRISPR spacer clusters were shared more widely among vent sites within Von Damm and Piccard compared to Axial Seamount, but microbial lineages were shared more widely among vent sites at Axial Seamount (Table 1).

**Table 1.** Distribution of vOTUs, CRISPR spacer clusters, and microbial hosts (classified at the lowest taxonomic level available based on 16S rRNA genes) across vent sites and vent fields at Piccard, Von Damm, and Axial Seamount. The Background and CTD1200 samples were excluded. Reads that mapped to 16S rRNA sequences that were classified as eukaryotes were excluded.

| Vent field | Type | Percent in more than one vent site | Percent in all vent sites in region |
| --- | --- | --- | --- |
| Von Damm | vOTUs | 13.71 | 0 |
| Von Damm | Spacer clusters | 5.1 | 0 |
| Von Damm | 16S rRNA host | 46.15 | 1.44 |
| Piccard | vOTUs | 10.78 | 0 |
| Piccard | Spacer clusters | 9.13 | 0 |
| Piccard | 16S rRNA host | 46.09 | 11.72 |
| Axial Seamount | vOTUs | 8.88 | 0.05 |
| Axial Seamount | Spacer clusters | 1.01 | 0 |
| Axial Seamount | 16S rRNA host | 54.55 | 2.36 |
| Type | Percent found in all three regions (Axial Seamount, Von Damm, and Piccard) |  |  |
| Viral OTUs | 0.02 |  |  |
| Spacer clusters | 0 |  |  |
| 16S rRNA host | 16.62 |  |  |

We created hierarchical dendrograms to assess the similarity of samples based on their viral content. Viral assemblages in samples from the Mid-Cayman Rise and Axial Seamount grouped separately (Figure 3A). Within Axial Seamount, samples taken in successive years from the same vent tended to have similar viral assemblage compositions (Figure 3A). In contrast, at the Mid-Cayman Rise, samples taken from the same site in two different years did not cluster together, and we observed weak clustering of samples by location (Figure 3A). Samples from Piccard vent field and Von Damm vent field did not cluster separately. Grouping of microbial communities based on classification of 16S rRNA reads in the metagenomes showed similar patterns to the viral assemblages. Samples collected from vents at Axial and the Mid-Cayman Rise grouped separately from each other, with only some clustering of samples at Piccard and Von Damm. Similarly, we observed stronger temporal clustering among samples at Axial Seamount than at Von Damm (Figure 3B).

**Figure 3.**
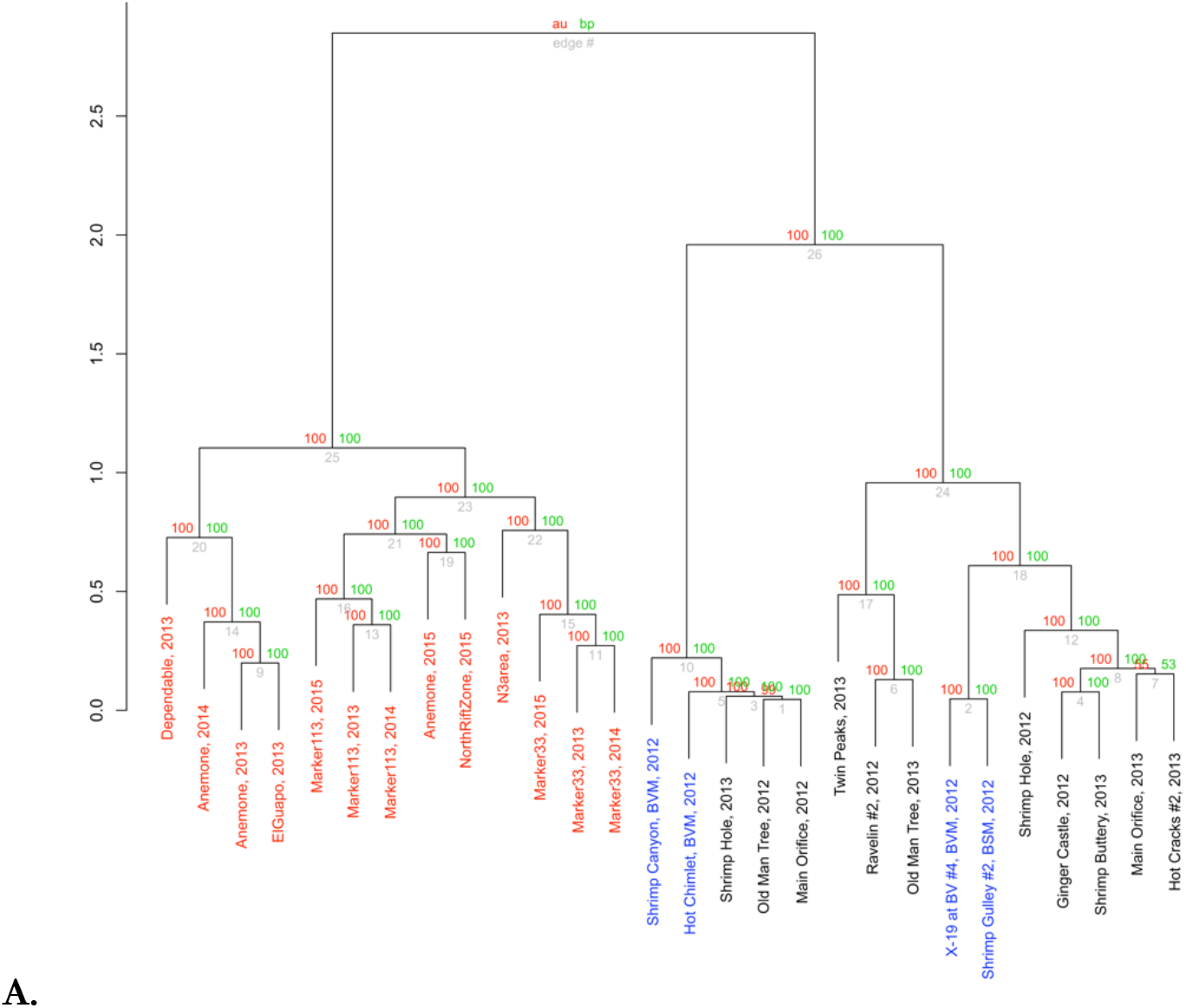
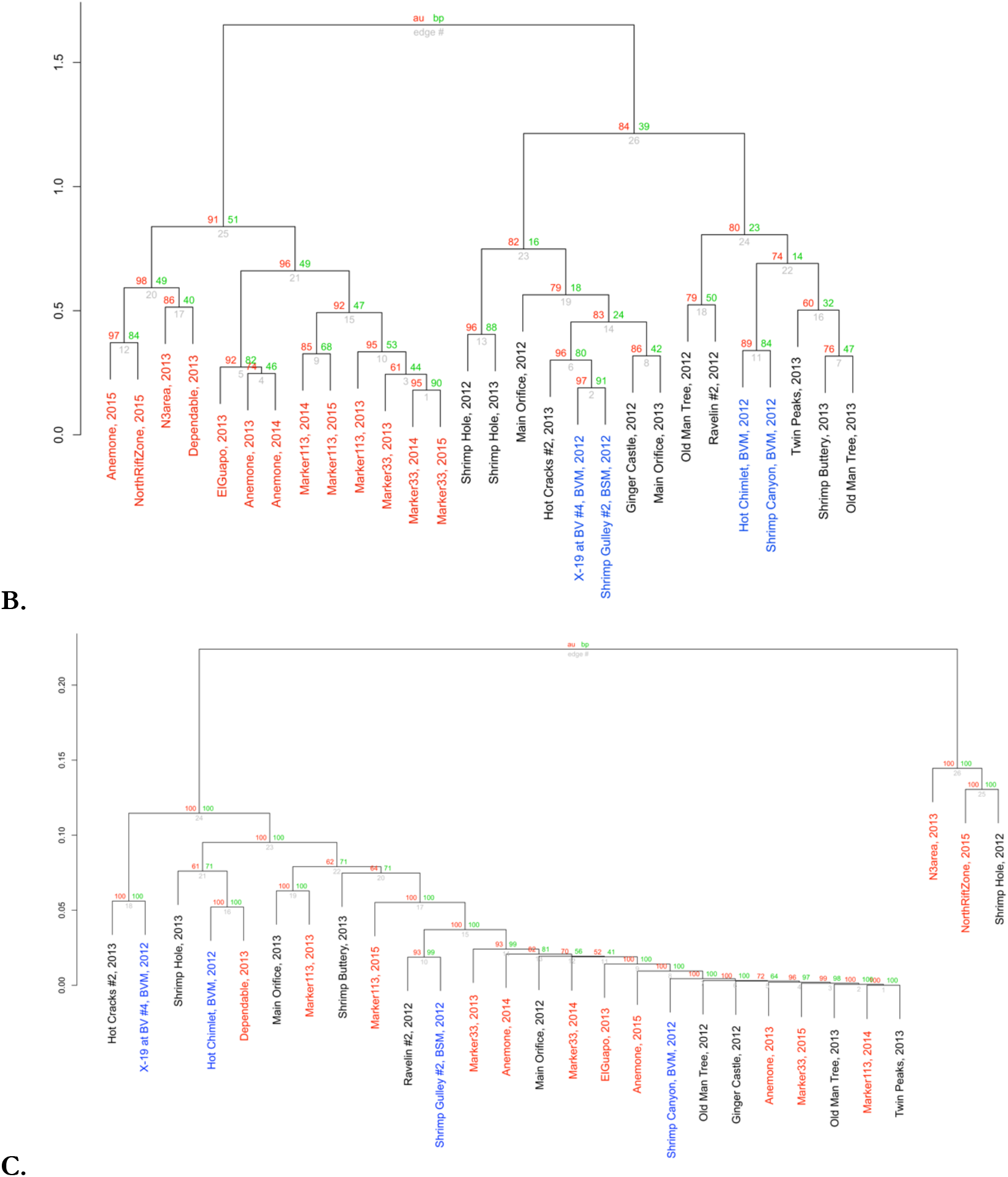
Hierarchical clustering of Von Damm (black) Piccard (blue) and Axial Seamount (red) diffuse flow samples based on A) viral assemblage, B) microbial host composition, and C) CRISPR spacer composition. For viral composition, the number of reads in each sample that mapped to each vOTU was calculated. For the host composition, the number of reads that mapped to 16S rRNA gene sequences of each host (classified at the lowest taxonomic level available) in each sample was calculated. Reads that mapped to 16S rRNA gene sequences that were either unclassified or classified as eukaryotes were excluded. For CRISPR composition, the number of CRISPR spacers in each sample in each spacer cluster was calculated. For all samples, the zero counts were replaced with estimates and a centered log-ratio (clr) transformation was applied. The y-axes indicate distance between samples as calculated by the ward.D2 method based on the transformed counts. Grey values at each edge are p-values (%). Red values are approximately unbiased (AU) p-values, computed by multiscale bootstrap resampling, and green values are bootstrap probability (BP) p-values, which are computed by normal bootstrap resampling. All uncertainty metrics in the dendrogram were calculated using the R package pvclust.

In contrast to the viral and microbial assemblages, hierarchical dendrograms based on CRISPR spacer compositions showed little clustering by location (Figure 3C). Based on spacer assemblages, samples from either Axial Seamount or the Mid-Cayman Rise did not cluster together, and samples taken from the same vent sites in different years did not cluster together. Very few CRISPR spacer clusters were found at multiple vent sites (Table 1).

### Networks of viral infection

Networks of viral infection generated from viral sequences, CRISPRs, and MAGs were used to examine the distribution and host specificity of vent viruses. These virus networks show connections between CRISPR spacers, which represent a record of previous viral infection in the host, and viral sequences recovered from the metagenome, which represent viral sequences present in the community at the time of sampling. Virus-host networks made for Axial Seamount (Figure 4A), Von Damm (Figure 4B), and Piccard (Figure 4C) indicate that vent viruses are restricted in terms of both host range and geographic distribution. We did not observe any virus-host connections shared between Piccard, Von Damm, or Axial Seamount. Within Axial Seamount, there were 89 vOTUs linked to 18 MAGs, and 105 connections between distinct pairs of vOTUs and MAGs (Figure 4A). We did not observe any connections between MAGs and vOTUs in any of the non-diffuse flow samples (Background, CTD1200 and the Anemone plume sample). The number of viral connections appeared to be related to, but was not significantly correlated with, the microbial hosts’ relative abundance (Supplementary Table 3): for example, a Clostridia MAG in the 2015 North Rift Zone sample (Clostridia_35) had high normalized coverage and was linked to nine vOTUs. We observed a high number of viruses shared among *Aquificae* MAGs sampled from the Anemone vent in 2013 and 2014 (Figure 4a). Finally, we observed only a single vOTU that was linked to MAGs of different taxonomic classes from different vent sites, with a vOTU connected to an *Ignavibacteria* MAG (Ignavibacteria_15) in the 2014 Marker 33 sample and an *Aquificae* MAG (Aquificae_43) in the 2013 Anemone sample (Figure 4A).

**Figure 4.**
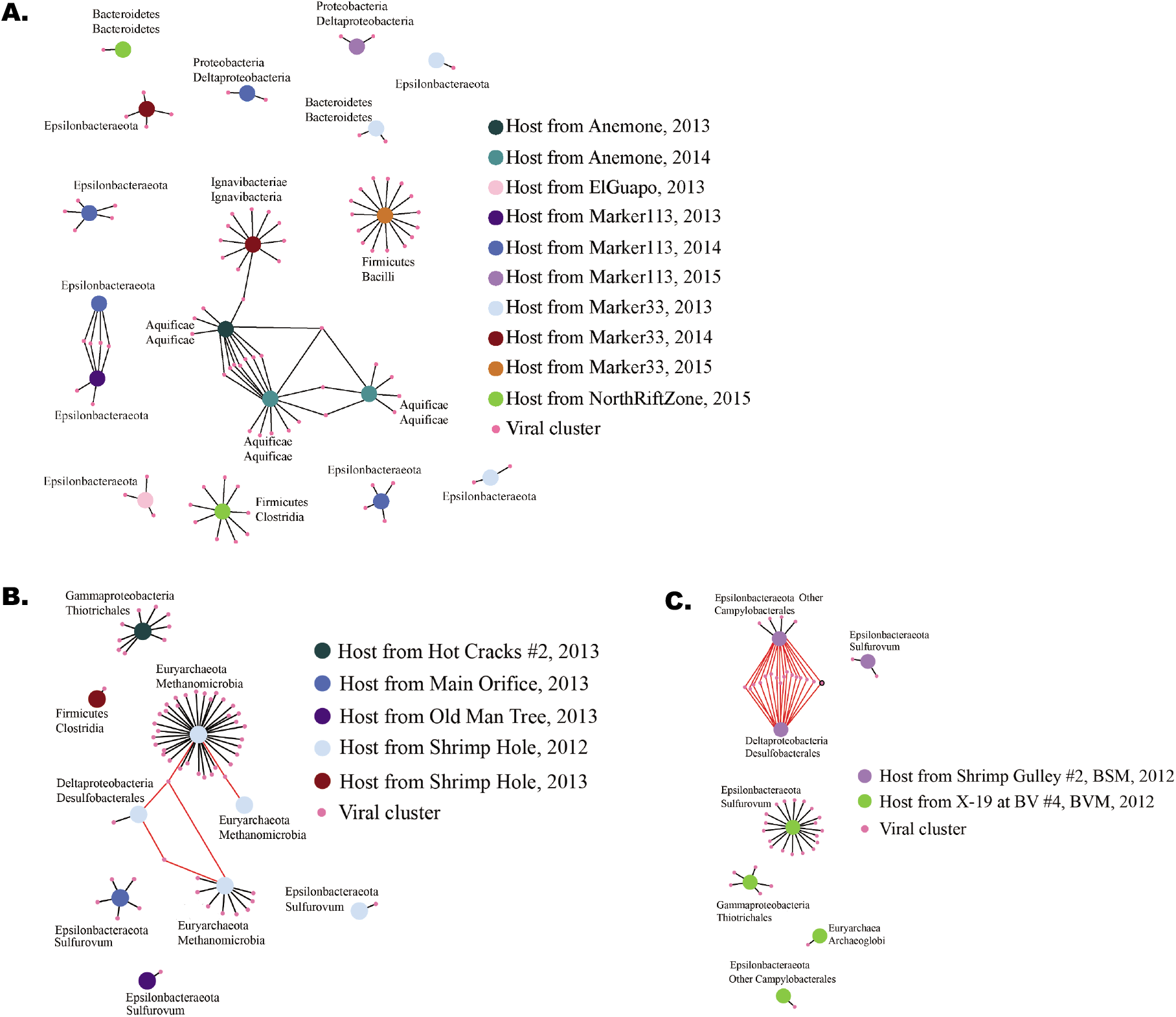
Infection network showing the links between viral clusters (vOTUs) and MAGs at A) Axial Seamount, B) Von Damm and C) Piccard. vOTUs were linked to MAGs via the spacers and direct repeats in CRISPR loci. Edges are colored red when a vOTU is linked to multiple MAGs in the same sample through the same direct repeat type. Due to the Crass algorithm, when the same direct repeat type is found in multiple MAGs in the same sample, it cannot be determined which MAG the spacers associated with direct repeat type came from. Therefore, the red edges are links between vOTUs and MAGs that could not be definitely proven. vOTUs with high relative abundance (top six most abundant) in at least one sample are outlined in black.

The virus infection networks reflected the geographic separation observed in the cluster dendrograms, and also revealed fairly narrow virus-host ranges for the viral sequences observed in our datasets. Within the Von Damm vent field in the Mid-Cayman Rise, there were 62 vOTUs linked to nine MAGs, with 66 connections between distinct pairs of vOTUs and MAGs (Figure 4B). The size of the viral infection network was similar in Piccard, where 48 vOTUs were linked to seven MAGs and there were 64 connections between distinct pairs of vOTUs and MAGs (Figure 4C). All of the vOTUs in the Von Damm and Piccard networks represented relatively rare viruses with the exception of one. This vOTU was present in seven of the Von Damm vents and all of the Piccard vents, was among the top six most abundant viruses in both the X-19 and Shrimp Gully #2 vents within Piccard, and was linked to a Campylobacterales and a Desulfobacterales MAG, both from Shrimp Gully #2. This was the only vOTU with high relative abundance in at least one sample that was present in the Axial Seamount, Von Damm, and Piccard viral infection networks (Figure 4). Some MAGs had more viral connections than others: for example, a *Methanomicrobia* MAG (Methanomicrobia_41) from the 2012 sample from Shrimp Hole within Von Damm was linked to 35 vOTUs (Figure 4B). However, this is not a direct indication of the number of different viruses infecting a specific strain because some MAGs had more CRISPR spacers than others, increasing the possibility of finding a viral connection. As with Axial Seamount, there were instances of highly abundant MAGs with many viral connections (Supplementary Table 3): a *Sulfurovum* MAG (Sulfurovum_99) in the 2012 sample from X-19 within Piccard had the highest normalized coverage across Von Damm and Piccard and was linked to 18 vOTUs, making it the MAG with the third highest number of viral connections within either network. Another *Sulfurovum* MAG (Sulfurovum_37) had a high relative abundance in the 2012 sample from Shrimp Gulley #2 within Piccard, and had two viral connections. Within Von Damm, Piccard, and Axial Seamount, we observed several cases in which viruses were connected to multiple microbial hosts within the network (Figure 4). However, in some cases the MAGs were linked to these shared vOTUs through the same CRISPR direct repeat type and were in the same sample, suggesting that these specific connections may have been due to matching CRISPR direct repeat types rather than true cross-infection. These have been indicated in Figure 4 (red lines).

Finally, an additional analysis of virus-host interaction was conducted by identifying prophages and auxiliary metabolic genes (AMGs) in MAGs. However, the fragmented nature of the assemblies and the absence of clear viral hallmark genes next to putative AMGs did not allow us to reach strong conclusions regarding the prevalence of lysogeny and AMGs in these hydrothermal vent samples. These analyses are described in the Supplementary Materials.

## Discussion

Viruses are important drivers of microbial mortality, ecology and evolution in the ocean, but studies of their distribution and impact in the deep sea are lacking. Our results indicate that viral diversity is high in venting fluids from hydrothermal systems, and the viruses we analyzed have restricted geographic distributions and host ranges. This implies that viruses do not spread widely between vent sites and that the viral role in mediating horizontal gene transfer across taxa and between vent sites is relatively restricted. However, the high abundance and diversity of viral sequences as well the large number of virus-host CRISPR connections implies rapid viral mutation and ongoing viral infection, indicating active and ongoing interactions between viruses and their microbial hosts in venting fluids from the hydrothermal vents examined here.

### Viral assemblages are taxonomically distinct and spatially restricted across vent sites

Previous studies of viruses in marine systems have observed viral populations to be commonly found across multiple samples (80), whereas others have found that most viruses are biogeographically restricted, with only a few cosmopolitan groups (81, 82). In hydrothermal vents at Von Damm, Piccard and Axial Seamount, we found high viral diversity with a limited distribution, potentially indicating rapid diversification in vents. Most vOTUs we observed were found only at individual vent sites, with very few cosmopolitan viruses. Previous work examining microbial distribution in hydrothermal systems through fine-scale 16S rRNA gene analyses indicates that while some microbial lineages are endemic to individual vent sites, others are widespread across vent fields (28, 40). Our work revealed that viral biogeographic patterns roughly corresponded to those of their microbial hosts. Taxonomic analysis revealed that there are distinct differences in viral taxonomy between vent sites--most notably, we observed a high abundance of *Myoviridae* at Axial Seamount that was not found at either Piccard or Von Damm. These observations indicate either restricted gene flow or strong environmental selection for specific groups within each site. Our analysis of viral sequence similarity using hierarchical clustering revealed similar results: there were some similarities between the viral assemblages in vents at Piccard and Von Damm, which are located approximately 20km apart in the Caribbean Sea, but the viral assemblages from Axial Seamount in the Pacific were very different from those at Piccard and Von Damm, despite the fact that vents at Piccard and Axial are both hosted in basalt and have metal-enriched, low pH fluids. Previous work on microbial community similarity and distribution in hydrothermal systems has shown that vents in close proximity are often more similar to one another in terms of microbial community structure than to geographically distant vents (4, 27, 40, 83). This may result from subseafloor plumbing that restricts fluid flux between sites, creating “islands” of microbial diversity that are distinct from one vent site to the next (26). Here, we show that these barriers to dispersal apply to viruses as well, and that viruses may be even more spatially restricted than their microbial hosts. Microbial lineages that spread between vent sites may face infection from novel viral strains not found in other vent sites. These endemic viral populations thus further shape distinct microbial community structure at individual vent sites.

Although the viral assemblages at hydrothermal vents had high diversity and restricted dispersal between vent sites and vent fields, our results show that the viral assemblages in vents persist over time, particularly at Axial Seamount where the same vents were sampled over a 3-year period. Viral assemblages from samples from the same vent sites at Axial across three years clustered together in the hierarchical dendrograms (Figure 3A), and our viral infection networks revealed that a number of viruses at Axial Seamount were linked to specific microbial hosts at the same vent over multiple years (Figure 4A). These patterns match those observed in the microbial communities (Figure 3B) and are consistent with previous work showing spatially restricted but temporally stable microbial communities over time at Axial Seamount (4, 27). Our work extends this to the viral world at Axial Seamount, indicating that viral assemblages follow the same temporal patterns as their microbial hosts, and that virus-host relationships persist over time. However, temporal stability in the viral and microbial community was generally not preserved at the Von Damm vent field for sites sampled in both 2012 and 2013. There were fewer sites and time points sampled at Von Damm compared to Axial Seamount, so it is unclear whether this results from a true biological signal or insufficient data.

### Vents host diverse and active assemblages of viruses with restricted host range

The viral infection networks show diverse microbial lineages across several samples infected by many different viruses at the time of sampling. Although individual microbial lineages could be targeted by multiple viral strains, the viruses we identified in these hydrothermal habitats had fairly restricted host ranges, infecting specific individual microbial strains. The only clear example of viral infection across microbial taxa emerged at Axial Seamount, where viral sequences associated with Ignavibacteria were also linked to Aquificales (Figure 4A). All other examples of viral infection across microbial taxa may either be true examples of viral-host cross-infection or may instead result from shared direct repeats. The limited host range of viruses from hydrothermal systems is consistent with previous work indicating that viruses tend to be host-specific (82), and we found very little evidence for generalist viruses, despite the fact that this has been reported previously (82, 84). Furthermore, we identified many viral connections to Epsilonbacteraeota, confirming that these highly successful microbial groups in hydrothermal vent diffuse flow fluids are susceptible to viral infection (3, 39, 85–87).

Matches between CRISPR loci and viruses provide an indication of which viruses are being targeted by the CRISPR immune response. We found that while some CRISPR spacers target relatively abundant viruses, most CRISPRs target relatively rare viruses in these systems. This is consistent with previous observations in an archaea-dominated hypersaline lake (88), where the vast majority of CRISPRs were found to target viruses with populations too small to allow for the assembly of contigs. The relative scarcity of viruses targeted by CRISPRs may result from an evolutionary arms race: CRISPRs limit the abundance of the viral populations they target, while concurrently, viruses undergo mutations, limiting the ability of CRISPR spacers to target them. Alternatively, these observations could result from an abundance of inactive spacers inherited over multiple generations. However, we do not believe this to be the case because CRISPR spacer clusters were infrequently present across multiple samples, suggesting that spacers were integrated on sub-generational timescales.

### Patterns recorded in microbial CRISPR loci do not reflect the contemporary viral assemblage

In contrast to the vOTUs, the CRISPR spacers did not demonstrate any clear biogeographic patterns. We expected CRISPR spacers to be more widespread than viruses since viral composition represents the virus community at the time of sampling, while spacers represent a history of viral infection. In contrast to our predictions, while both had limited distributions, we found viruses to be more widespread than CRISPR spacers at all scales examined (Table 1). These results are in contrast to previous studies of CRISPR spacer biogeography in terrestrial hot springs, where both viral and CRISPR spacers showed clear biogeographic structure (89). This suggests that there is selective pressure for CRISPR spacer composition to evolve more rapidly than viral sequences via the loss or mutation of CRISPR spacers. However, it is also possible that we found viruses to be more widespread because of undersampling of CRISPR spacers, or our use of different clustering algorithms for viruses and spacers: spacers may have been clustered at a finer resolution, resulting in a narrower distribution for each spacer.

Given that CRISPR spacers provide a history of infection, comparing the record of past viral infections via CRISPR arrays with viral sequences in the metagenomes can provide insights into whether CRISPR arrays provide an accurate representation of the viral assemblage at the time of sampling, as well as the rate at which CRISPR spacers are accumulated. We did not find that the most common or cosmopolitan viruses and CRISPR spacers matched each other. This may arise from a temporal mismatch: it takes time for spacers to be incorporated into and lost from CRISPR loci as virus abundances shift and viruses evolve. Additionally, just one SNP (which we allowed for when aligning spacers to viruses) can prevent a CRISPR spacer from providing resistance against a virus (66). This could explain the discrepancy between viruses and spacers: once resistance of a spacer to a particular virus is suppressed, the population of the virus is freed to shift independently from the spacer in the host population. The discrepancy we observed between viruses and spacers is important to note when attempting to use CRISPR spacers to study viral populations or vice versa.

Examination of the prophage abundance in MAGs revealed a relatively high abundance of prophage encoded in each MAG, confirming previous results indicating that lysogeny is a common lifestyle in hydrothermal systems (14, 15). However, while examination of the gene content of viral contigs revealed several ORFs encoding a range of functions including cell membrane function and energy production, it was difficult to prove conclusively that these genes were viral-encoded AMGs rather than potential cellular contamination, and thus no concrete conclusions were provided here (see Supplementary Materials). Previous research has suggested that vent viruses encode AMGs (18, 19), and thus it is likely that many of these genes represent viral-encoded AMGs, but further research is necessary to determine AMG diversity and prevalence across vent systems.

Taken together, our results show that hydrothermal vent viruses are active, abundant, and diverse. These viruses are restricted in their host range and biogeographic extent, but their interactions with hosts persist over time. Thus, while viruses in venting fluids from deep-sea hydrothermal systems have the capacity to play an important role in driving the evolution and ecology of microbial communities, their influence appears to be highly localized to specific regions and taxa. Future work examining viral diversity and distribution across higher resolution transects over space and time should reveal further insights into the extent of the viral influence in deep-sea hydrothermal vents.

## Supporting information

all Supplementary Tables

## Acknowledgements

We thank Julie Reveillaud, Caroline Fortunato, and Emily Reddington for support in the collection and generation of metagenomic data, and Chip Breier, David Butterfield, Bill Chadwick, Chris German, Jim Holden, Jill McDermott, and Jeff Seewald for sample collection support at sea. Funding for ET was provided by Carleton College. REA was supported by a NASA Postdoctoral Fellowship with the NASA Astrobiology Institute. This work was supported by a NASA Exobiology grant number 80NSSC18K1076 to REA and JAH; a NASA Astrobiology Science and Technology for Exploring Planets (ASTEP) grant NNX-327 09AB75G and a grant from Deep Carbon Observatory’s Deep Life Initiative to JAH; the NSF Science and Technology Center for Dark Energy Biosphere Investigations (C-DEBI) to JAH; and the Gordon and Betty Moore Foundation Grant GBMF3297 to JAH. Samples were collected from the Mid-Cayman Rise with the assistance of the captains and crew of the R/V Atlantis and R/V Falkor as well as ROVs Jason and Nereus. For Mid-Cayman Rise, ship and vehicle time in 2012 were supported by the NSF-OCE great OCE-1061863 to Chris German and Jeff Seewald and in 2013 by the Schmidt Ocean Institute during cruise FX008-2013 aboard the R/V Falkor. Samples collected from Axial Seamount were collected with the assistance of the captains and crew of the R/V Falkor, R/V Thompson, and R/V Brown as well as the ROV ROPOS and Jason groups, and in 2013 the Schmidt Ocean Institute during cruise FK010-2013 aboard the R/V Falkor. This is C-DEBI Contribution number [number to be determined].

## SUPPLEMENTARY MATERIAL

### Viruses and prophages in MAGS and viral and prophage genes

In order to identify viral genes and putative auxiliary metabolic genes (AMGs), VirSorter (69) v1.0.3 was used to extract putative viral and prophage sequences from metagenomic assemblies and from individual MAGs. In the case of identifying putative auxiliary metabolic genes, we chose to maximize accuracy rather than recovery. Therefore, we used VirSorter for prophage and AMG identification because VirSorter relies on known viral hallmark genes for viral identification and thus is a more conservative tool for positive viral identification. Category one, category two, category four (prophage), and category five (prophage) VirSorter sequences were used in analyses. Putative viral and prophage sequences from all assembled contigs were annotated via Prokka (69, 70) v1.14 with the taxonomic identifier set to “bacteria” as well as to “viruses.” We assigned categories to the ORFs that had been identified with the “bacteria” taxonomic identifier using the Clusters of Orthologous Groups of proteins (COG) database (71, 72). We conducted a second round of annotation for ORFs identified by Prokka using the “viruses” taxonomic identifier using VirSorter; we classified these ORFs as having viral function if they were annotated with terms that included “capsid,” “tail,” “spike,” “terminase large subunit,” “portal,” and “coat.”

Of the 74 Mid-Cayman Rise MAGs, 28 (38%) had at least one putative prophage sequence counting VirSorter categories one, two, four, or five, and 6 (8%) had a putative prophage counting only VirSorter categories four and five (Supplementary Table 4). Of the 98 Axial Seamount MAGs, 41 (42%) had at least one putative viral or prophage sequence counting VirSorter categories one, two, four, or five, and 6 (6%) had a putative prophage sequence counting VirSorter categories four and five (Supplementary Table 4). Given the uncertainties associated with binning viral contigs into MAGs, the true number of prophage in hydrothermal vent microbial genomes is likely between these endmember values. VirSorter has not been verified in hydrothermal systems; however, it is commonly used in other non-surface ocean systems (61, 84, 85).

To determine whether viruses and prophage in these samples carried auxiliary metabolic genes (AMGs), we annotated and characterized viral and prophage gene content in the viral sequences recovered from the metagenomes. The vast majority of genes within the putative viral and prophage sequences were annotated as belonging to the COG categories replication and repair; nucleotide metabolism and transport; and post-translational modification, protein turnover, chaperone functions. Many genes were also annotated as belonging to the cell wall/membrane/envelope biogenesis category. Within the Mid-Cayman Rise, a total of 106 ORFs within putative viral or prophage sequences identified in 10 different metagenomes fell within the broad COG category of “metabolism” (Supplementary Fig. 5). This included genes categorized as energy production and conversion, amino acid transport and metabolism, nucleotide transport and metabolism, carbohydrate transport and metabolism, coenzyme transport and metabolism, lipid transport and metabolism, and inorganic ion transport and metabolism. Samples taken from X-19 and Shrimp Hole (2012) had a high number of metabolism genes (30 and 28, respectively) on putative viral or prophage sequences compared to the other Mid-Cayman Rise samples. A higher proportion of genes on putative viral or prophage sequences were related to metabolism at the Mid-Cayman Rise than Axial Seamount (Supplementary Fig. 5). At Axial Seamount, we identified a total of 185 ORFs on putative viral or prophage sequences from 14 different metagenomes that fell within the broad COG category of “metabolism.” However, it is important to note that none of the observed metabolism genes were surrounded by confirmed virus genes (i.e., known viral ORFs both upstream and downstream of the metabolic ORF), and therefore we cannot rule out the possibility that some of these genes were microbial in origin.

### Auxiliary metabolic genes and the role of virus-driven horizontal gene transfer in these systems

Previous work has suggested that viruses in hydrothermal systems have the genomic capacity to alter their hosts’ microbial metabolism through harboring auxiliary metabolic genes (AMGs) (17–19). We found a higher percentage of microbial genomes with putative viral sequences (40%) than has been previously reported for single-cell genomes (SAGs, 10%) in diffuse flow hydrothermal fluids (15). The high abundance of prophage identified here is consistent with previous work documenting a high incidence of prophage in diffuse flow systems compared to deep seawater (14).

The ORFs on the viral contigs identified in these hydrothermal vent metagenomes encoded a wide range of functions, some of which may function as auxiliary metabolic genes. Many viral contigs contained genes related to cell wall or membrane proteins. The function of these genes is unknown; it is possible that these genes are involved in the synthesis of membranes surrounding viral capsids. Genes related to outer membrane proteins and protein glycosylation are commonly observed as variable genes within microbial pangenomes (94–100), possibly as a means to vary membrane proteins to evade viral infection. Viruses often act as a source of genes for horizontal gene transfer via transduction, and it is possible that the introduction of these novel genes may enable some microbial strains to avoid infection by other viruses. We also observed several genes with functions related to energy metabolism, inorganic ion transport, and signal transduction. Previous work has found that genes related to inorganic ion transport are differentially distributed in the *Sulfurovum* pangenome according to nutrient availability (39) and further work on viral sequences across vent sites is needed to determine whether these genes are carried by viruses to benefit their hosts. Moreover, work on viruses in hydrothermal plumes (19) and in diffuse flow vent fluid (18) has indicated that viruses in these systems encode energy-metabolizing AMGs, potentially to supplement the host’s ability to generate sufficient energy for cellular processes during the course of infection. The abundance and diversity of metabolic genes observed on these contigs provide further evidence that viral-encoded AMGs are widespread and diverse within hydrothermal vent ecosystems. These genes also have the potential to be horizontally transferred via viral transduction. However, our analyses of viral biogeography in hydrothermal vents has revealed virus-host interactions to have limited distributions because viruses are spatially restricted and highly host-specific. Thus, the potential for viruses to act as mechanisms for horizontal gene transfer between spatially and phylogenetically distant hosts is likely limited.

## SUPPLEMENTARY FIGURES

**Supplementary Figure 1.**
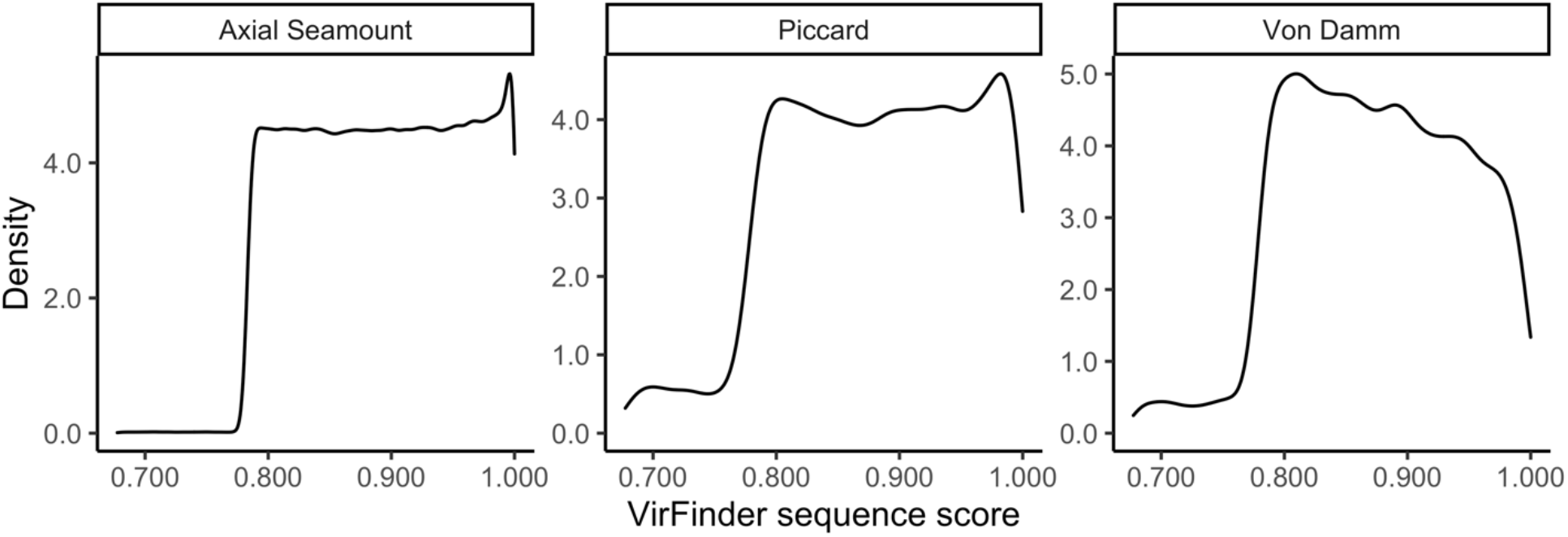
Distributions of the VirFinder score of putatively viral sequences identified at the Axial Seamount, Piccard, and Von Damm vent fields. The majority of VirFinder sequences are assigned scores above 0.8 at all three vent fields.

**Supplementary Figure 2.**
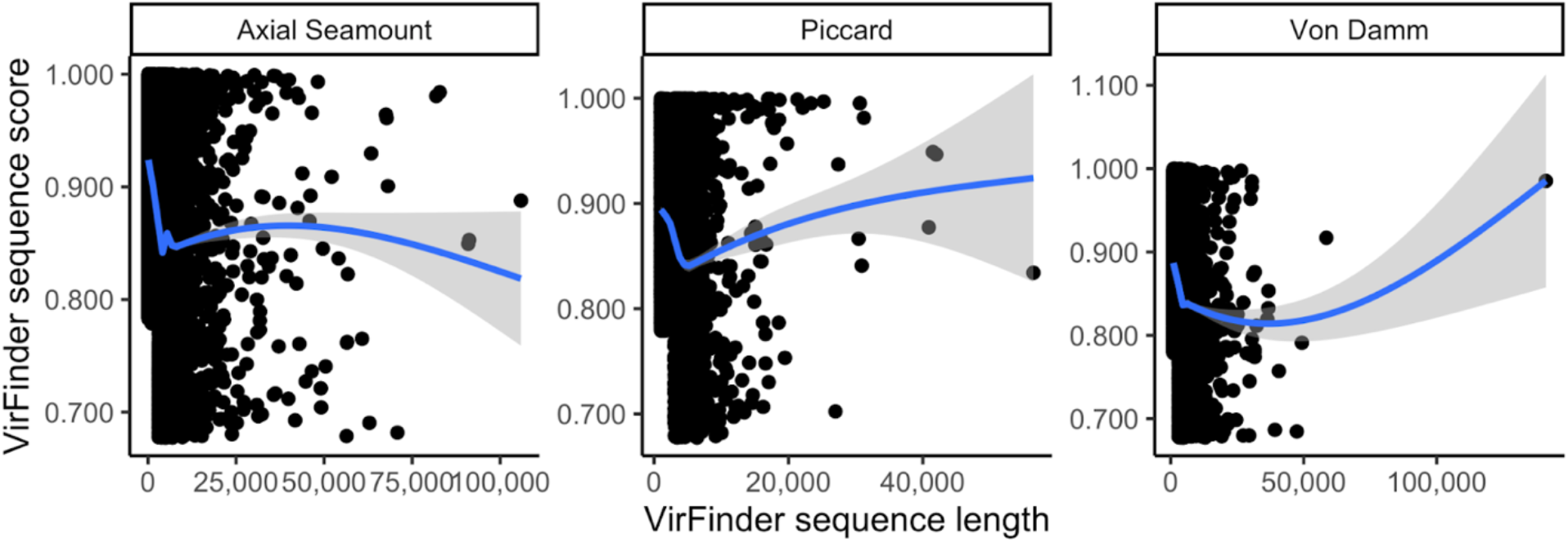
VirFinder sequence score vs. length at the Axial Seamount, Piccard, and Von Damm vent fields. The VirFinder score does not increase with sequence length for the majority of sequences.

**Supplementary Figure 3.**
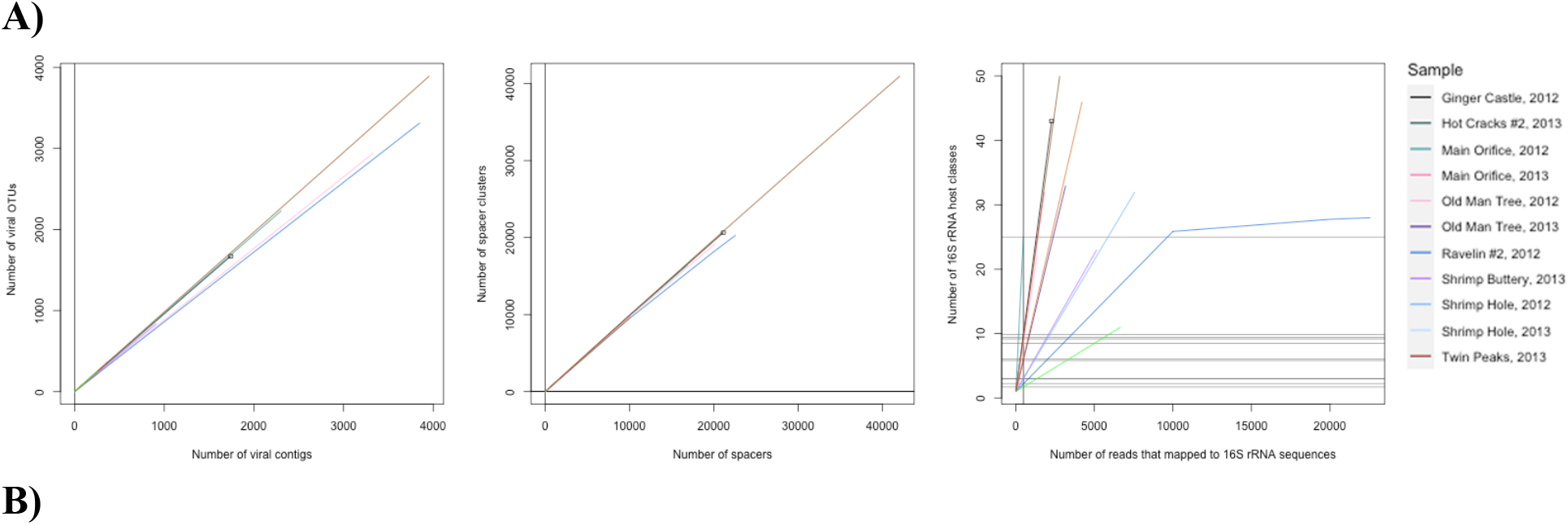
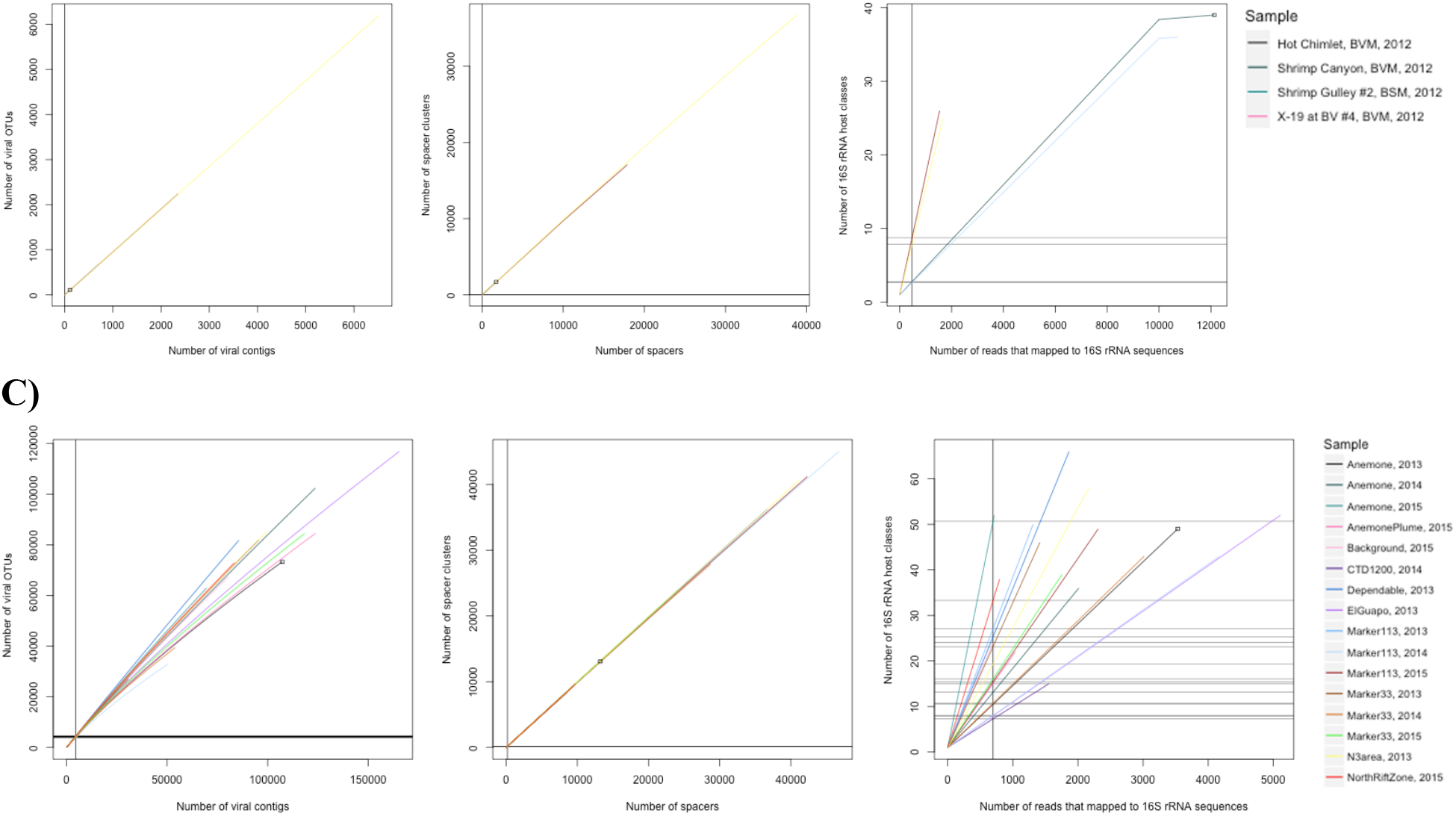
Rarefaction curves for viral OTUs, spacer clusters, and 16S rRNA host classes from samples collected at A) Von Damm, B) Piccard and C) Axial Seamount. There is no viral rarefaction curve for Old Man Tree, 2012 because there were no viral OTUs in this sample.

**Supplementary Figure 4.**
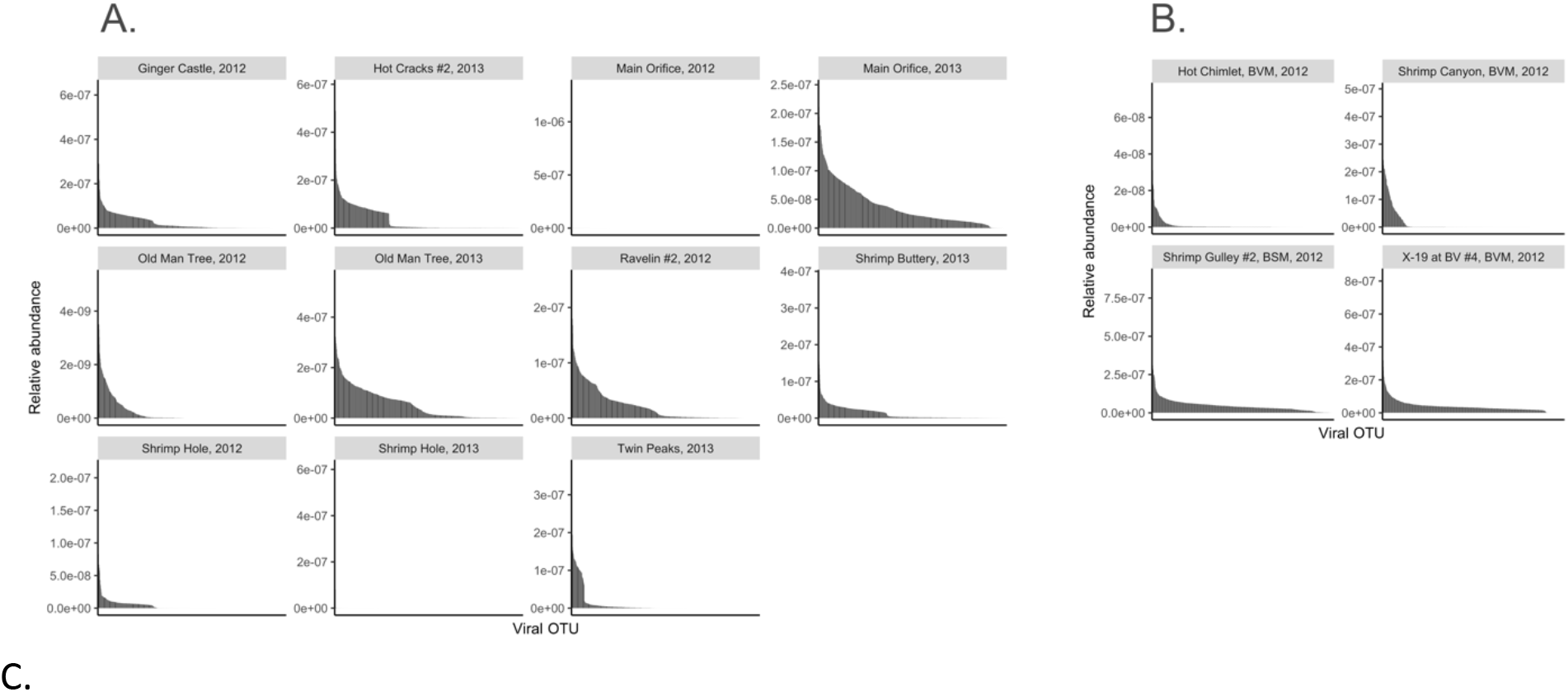
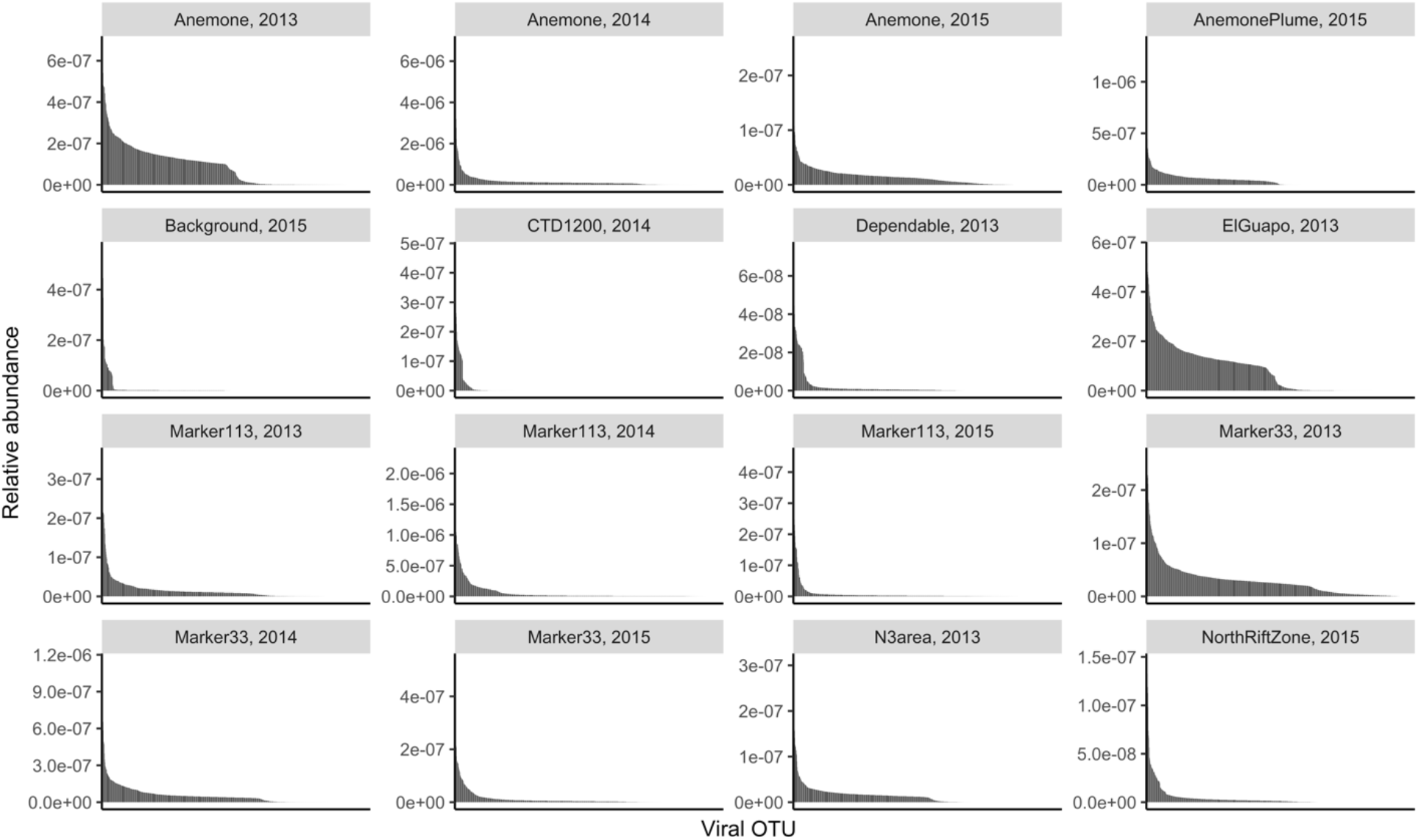
Rank abundance curves of vOTUs at A) Von Damm, B) Piccard and C) Axial Seamount. The y-axis represents the number of reads that mapped to each vOTU divided by the total length of the viral contigs in the cluster and the number of reads in the sample. The x-axis represents the vOTUs ordered according to relative abundance, with the most abundant vOTUs at the left. For each vent field, the 1000 most abundant viral OTUs are shown.

**Supplementary Figure 5.**
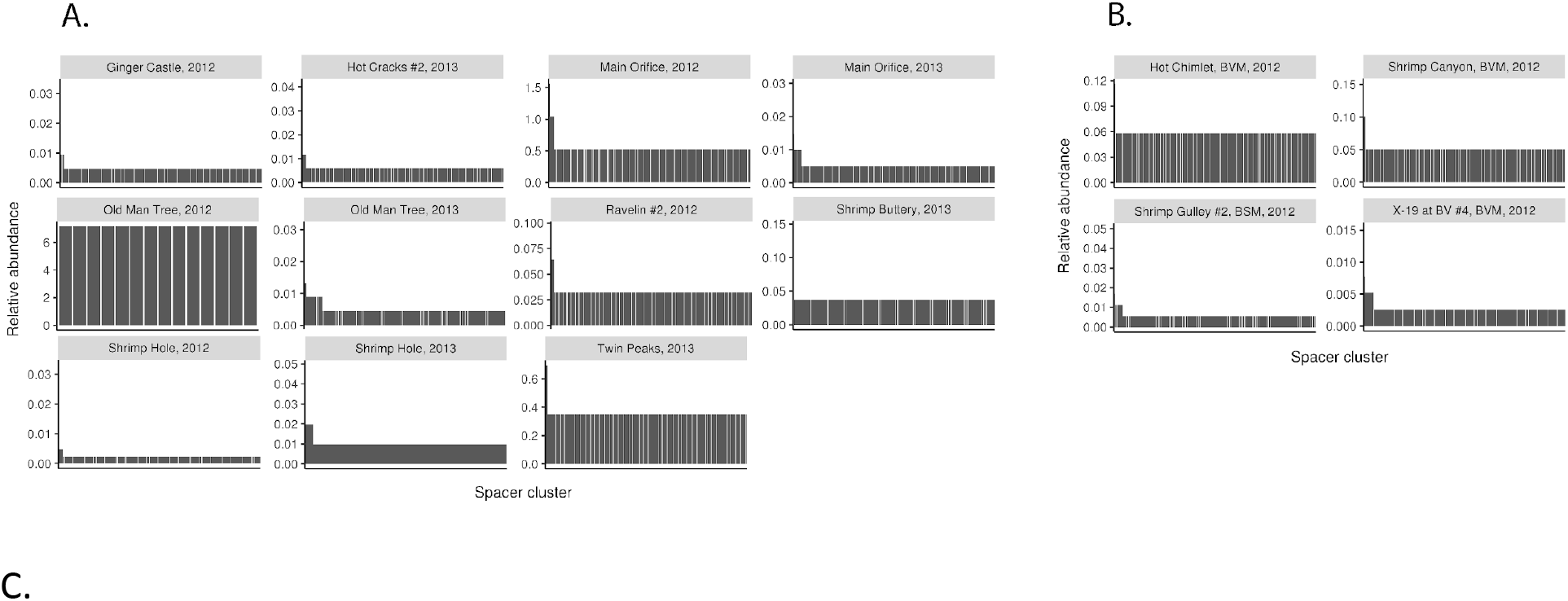
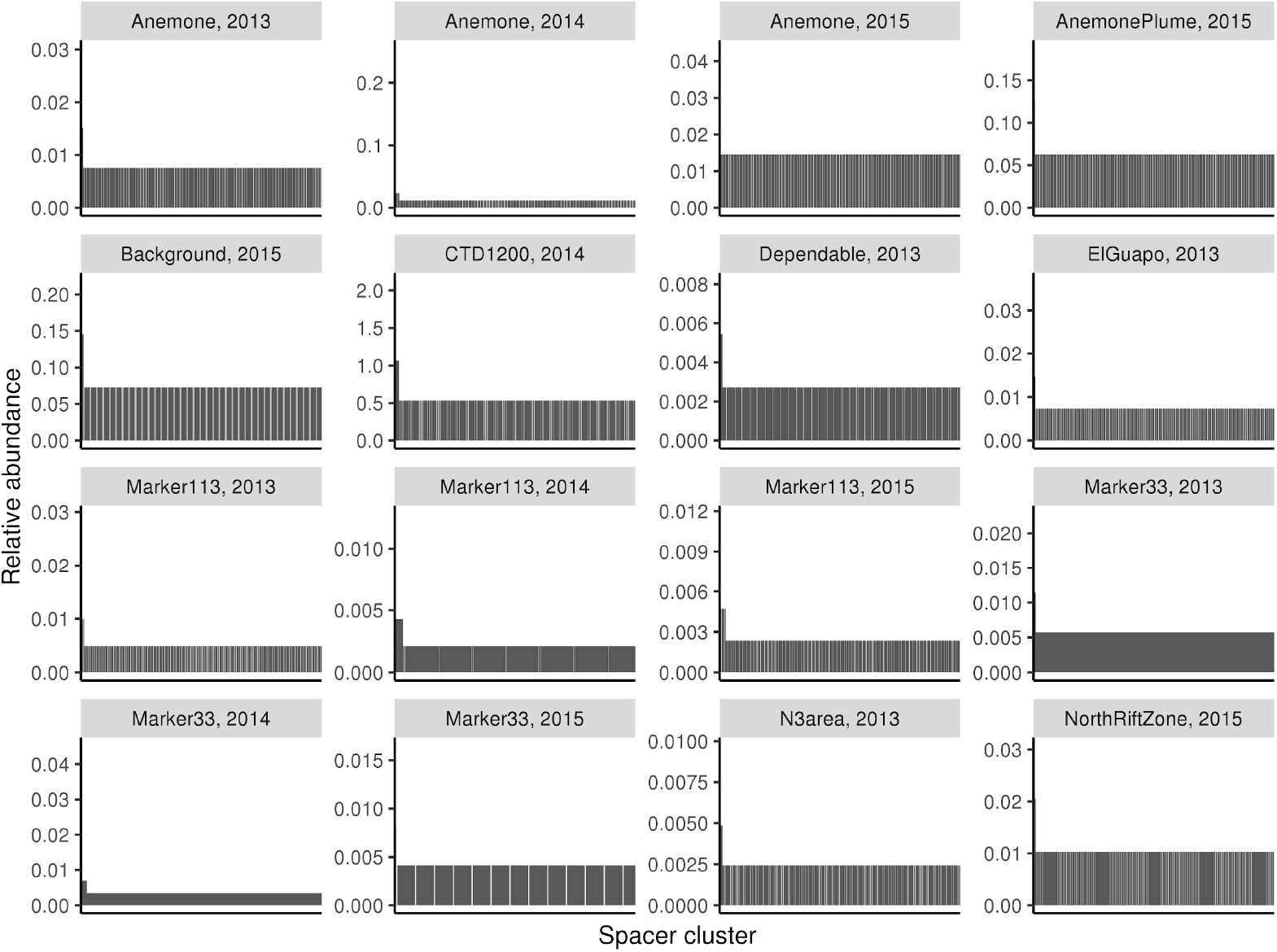
Rank abundance curves of CRISPR spacer clusters in A) Von Damm, B) Piccard and C) Axial Seamount. The y-axis represents the percent of all CRISPR spacers in a sample falling into that cluster. The x-axis represents the CRISPR spacer clusters ordered according to relative abundance, with the most abundant clusters at the left.

**Supplementary Figure 6.**
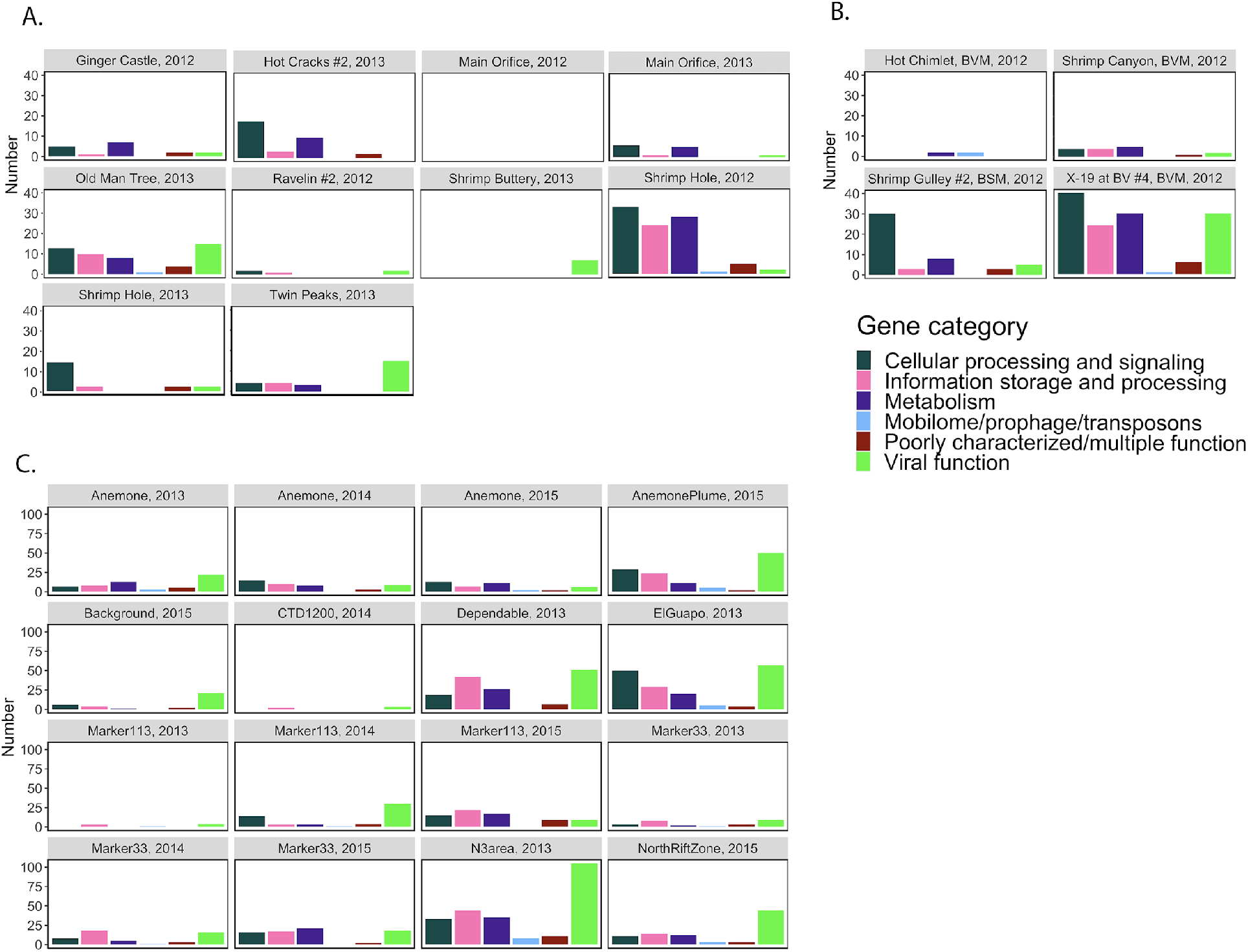
COG categories assigned to ORFs on contigs identified as being derived from viruses or prophage by VirSorter. A) Samples from Von Damm; B) Samples from Piccard; C) Samples from Axial Seamount.

**Supplementary Tables are provided in Excel format.**

## Notes

### Competing Interest Statement

The authors have declared no competing interest.

### Summary of Updates

This revision includes some minor edits to the text (including redefining viral clusters as "vOTUs") as well as the figures.

